# A quantitative molecular model of the lymphocyte death timer

**DOI:** 10.1101/2023.10.25.563681

**Authors:** Michelle Ruhle, Evan Thomas, Edward Dann, Nicole Gottscheber, Charis E. Teh, Daniel H.D. Gray, Mark R. Dowling, Susanne Heinzel, Philip D. Hodgkin

## Abstract

When stimulated, individual lymphocytes program times for division and death that are inherited within families, revealing a common timing mechanism transmitted over generations. Here we describe a threshold-based mechanism for the time to die. By comparing protein levels in control and apoptosis disabled B and T lymphocytes, we show that death can be predicted by a cooperating ensemble of BCL-2 family proteins falling below a critical threshold. Single cell measurements predict the time of death with a simple formula, where additional inhibition factors explain accelerated death induced by BH3 mimetic compounds. Thus, we identify the death timer as a protein-threshold device that underlies signal integration machinery. Together these results reveal that predicting lymphocyte behavior at single cell level, in complex environments, is possible with modular models that incorporate independent timers and heritability features of critical proteins.

## MAIN TEXT

The effector cells of the adaptive immune response, T and B lymphocytes, typically circulate in a resting, quiescent state. When stimulated by foreign antigen, along with other costimulatory ‘danger’ signals, T and B cells program a burst of proliferation after which, they appear to automatically re-enter a quiescent state and many die, leaving behind a set of long-lived memory cells ^1, 2^. Quantitative analysis of lymphocyte dynamics under controlled stimulation conditions in vitro reveals similar behavior: Cells undergo a proliferative burst that varies in proportion to stimulation strength that is followed by death, enabling careful dissection of the control features ^3, 4^. Such studies indicate that both T and B cells can adapt their survival and proliferation times independently, effectively placing alternative outcomes in competition within cells. Further, within a single resting cell, stimulation induces cellular fate changes that can be delayed in time and heritable in clonal families leading to all descendant cells dividing the same number of times before stopping (their division destiny) ^5, 6, 7, 8, 9^ and dying in a similar generation ^5, 10^. The regulation of both division destiny and the times to die are ‘tunable’ in clones as they vary with strength of stimulation and combinations of signals ^11, 12^. These striking dynamic features raise the question of how molecular controls over division and death operate and are tuned to regulate the scale and rate of clearance of an immune response.

The mechanism for timed regulation of division progression is known. Heinzel et al. identified the oncoprotein Myc as the key driver of division progression ^12^. This study found stimulation of T and B cells induces Myc in proportion to signal strength and over time the production rate wanes, the protein level diminishes and, when below an identifiable threshold, cells stop dividing. The kinetics of the rise and fall in Myc level is unaffected by cell division explaining the strong relatedness in fate by clonal family members ^5, 12^. Together these features identify the control of division progression as a heritable cellular ‘timer’ ^4, 13^. Importantly, control of division progression by Myc and control of cell death were separable; division destiny was reached at an identical time if cell division was slowed down, or if cell death was prevented ^12^. These heritable timer features inform a mathematical model of cell dynamics, Cyton2, that fits T and B cell proliferation and survival data ^11^. Applying this model to data in Marchingo et al. ^13^ revealed that times to division destiny and times to die were independently ‘added’ by combining stimulatory signals ^11^.

How might a tunable cell death timer operate? By analogy with Myc, we postulated a mechanism where critical proteins change expression over time and cells die when levels fall below a threshold. Components for such a mechanism are well described. BCL-2 protein family members can vary in expression over time and combine to set thresholds for initiating apoptosis. Broadly speaking, members of this protein family can be separated into three factions: the apoptosis-initiating BH3-only proteins, the pro-survival proteins and the apoptotic effector proteins. The activation of the effector proteins, BAX and BAK, is a critical step in the apoptotic cascade, resulting in permeabilization of the mitochondrial membrane^14, 15, 16^. In viable cells, the activation of BAX and BAK is restrained by the binding of pro-survival molecules including BCL-2, BCL-xL, MCL-1 and A1 ^17, 18, 19^. Pro-apoptotic BH3-only proteins, such as BIM, NOXA and PUMA are upregulated in response to cellular stress and other signals. BH3-only proteins bind to pro-survival BCL-2 family members, thus alleviating their inhibition of BAX and BAK, and can also directly activate the effector proteins to induce apoptosis ^14, 18, 20, 21, 22, 23, 24^. Evidence suggests multiple pro-survival BCL-2 proteins expressed in the same cell can cooperate to neutralize pro-apoptotic proteins and regulate maintenance of cell survival ^25^. Further, similar cells exhibit broad, non-genetic differences in expression level of these molecular components, rendering each cell unique in behavior and responsiveness to stimulation or drug sensitivity ^26, 27, 28^. Thus, the death timer mechanism is potentially complex, requiring a summation of pro- and anti-apoptotic proteins in individual cells to determine whether a threshold is breached.

The role for apoptosis in both development and tissue homeostasis has led to wide acceptance that cells are intrinsically programmed to die, and that cell survival depends on the continuous repression of this default program ^29, 30, 31, 32^. Consistent with this idea, Mason et al, proposed that platelet lifespan was determined by a ‘‘molecular clock’’, with the loss of pro-survival protein BCL-xL as platelets age sufficient to trigger Bak mediated apoptosis ^33^. Furthermore, studies of thymocytes and cancer cells show they vary in their level of pro- and anti-survival proteins and therefore are more or less sensitive to BH3 mimetic induced cell death^34, 35, 36^. This concept of ‘apoptotic priming’ where each cell can be close to, or distant from, an apoptotic threshold is suggestive of a quantitative relationship that may be tractable with a protein level model. Here, we build on these concepts to search for quantitative mechanisms controlling the tunable death timer in lymphocytes.

## RESULTS

### BCL-2 family proteins quantitatively regulate the timing of B cell death following stimulation

CpG activation of naïve B lymphocytes by engagement of TLR9 is a well-studied and tractable system for tracing the connection between proliferation and cell death ^5, 12, 37^ ^38^. The system is highly quantitative and the mean times and variances for division entry, subsequent division rate, division destiny and death can be determined using time series division tracking data fitted to the Cyton2 model ^11^. The variation in times for each fate are empirically well fitted by lognormal probability distributions ^5, 11, 39^. As death of CpG activated B cells is a heritable, timed process, we sought the underlying molecular mechanism of control in this system. We hypothesized that dynamic changes in the expression level of key BCL-2 protein family members would control the timing of apoptosis in each cell (Fig. 1a). If correct, modulating the levels of these proteins or inhibiting their action should quantitatively alter the average time to die. To test this prediction, and identify the key players in this system, we manipulated the effective levels of BCL-2 family proteins using genetic ablation or treatment with specific inhibitors, termed BH3 mimetics. Naïve B cells were isolated, CTV labelled, stimulated with CpG and sampled over time as previously described ^13, 40, 41^. B cell survival was monitored after BIM deletion ^42^ or treatment with MCL-1 inhibitor (MCL-1i), S63845 ^43^, BCL-2 inhibitor (BCL-2i), ABT199 ^44^ and BCL-xL inhibitor (BCL-xLi), A1331852 ^45^ 24 hours after activation.

**Figure 1.**
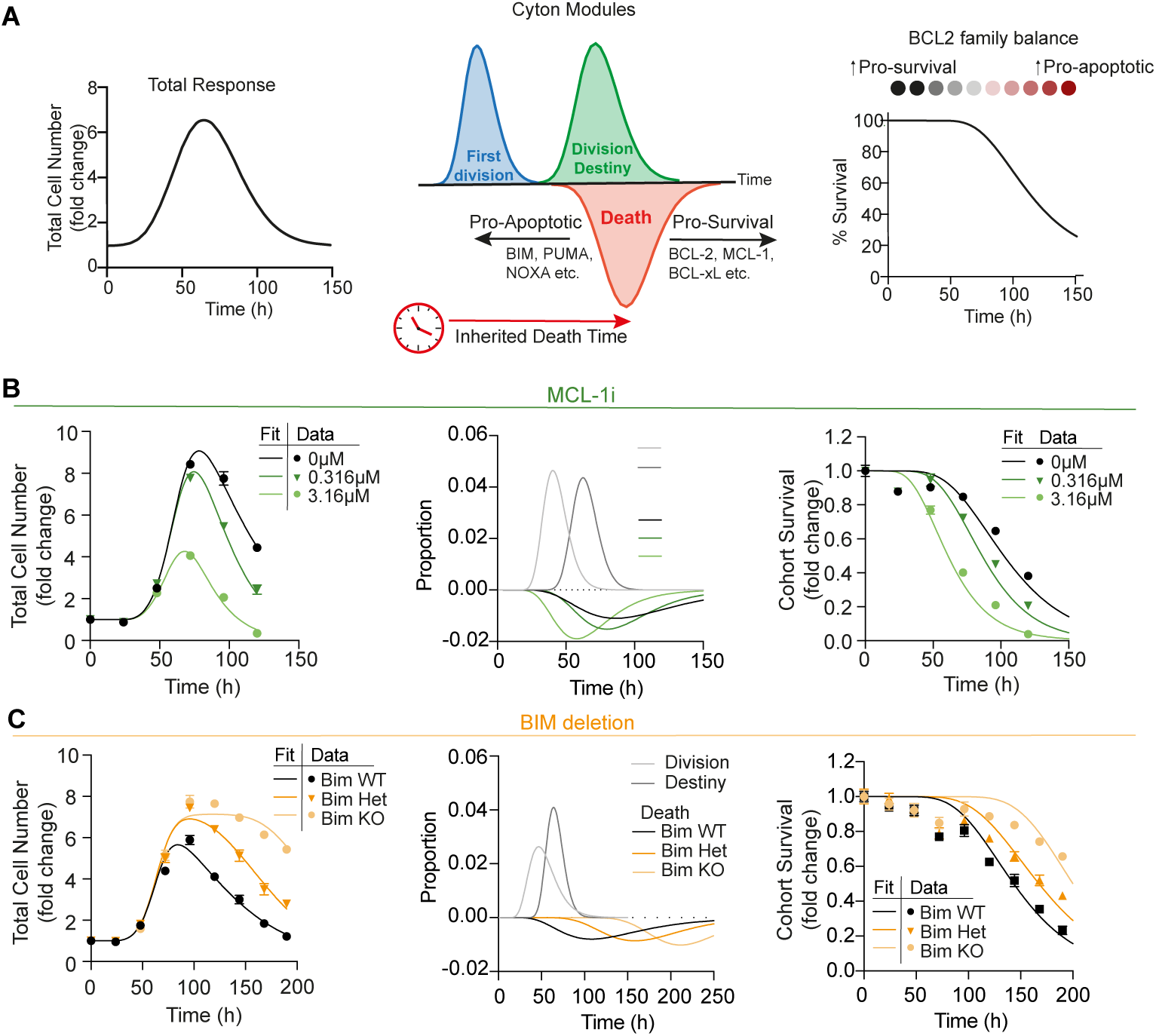
B cell death times are quantitatively regulated by levels of multiple BCL-2 family proteins. (**a**) Hypothesis of death time regulation by BCL-2 family proteins. (**b-c**) Analysis of CpG activated B cells following (**b**) MCL-1 inhibition (S63845) or (**c)** genetic deletion of BCL-2l11 (BIM). Total cell numbers and cohort survival over time were quantified. Using Cyton2 modeling, Cyton parameter distributions were determined and model fits of total cell number and cohort survival over time were plotted alongside the measured data. Measured data represented as mean ± SEM. Data representative of three (b) or two (c) independent experiments.

Using the 7-parameter Cyton2 model ^11^, we fit time series data from B cells stimulated with CpG alone or together with BH3 mimetic treatment. As the BH3 mimetics tested did not alter B cell proliferation (Supplementary Fig. 1a), the division parameters, including time to first division, subsequent division times and division destiny times determined for untreated CpG cells were applied across all conditions, reducing the number of free parameters. Cyton2 fitted values of live cell number over time and the effects on death time distributions were determined. The fitted time to die distribution can also be determined directly by plotting precursor cohorts ^41^ and fitting a lognormal survival curve, avoiding the need to assume Cyton2 principles as the controlling function (Supplementary Fig. 1b). In response to MCL-1i (Fig. 1b), a dose dependent reduction in the time to die distribution and cohort survival was observed. Similar effects on survival and time to die distributions were observed following BCL2i and BCL-xLi (Supplementary Fig. 1c-d). Single drug effects were further compounded when BH3 mimetics were used in combination, with paired or triple BH3 mimetic treatment reducing B cell numbers and shifting death time distributions markedly, indicating that these three proteins are collectively playing a major role in the survival of these cells (Supplementary Fig. 1e).

When the same analysis was applied to CpG stimulated B cells with partial or complete deletion of the pro-apoptotic protein BIM, Cyton2 modelling was able to resolve extended death time distributions, reflective of BCL-2l11 (BIM) gene dosage. The heightened total cell numbers over time were a function of quantitative increases in cell survival (Fig. 1c). Together, these results support the death timer hypothesis, with average death times shifted up or down in response to changing the effective cellular concentrations of both pro-survival and pro-apoptotic BCL-2 family proteins respectively.

### Preventing apoptosis unmasks censored protein expression patterns, indicating expression thresholds for survival maintenance

As the time to die mechanism of CpG activated B cells relied on the availability of BCL-2, MCL-1, BCL-xL and BIM, we investigated the expression of these molecules over time. To enable resolution of these parameters uncensored by cell death, we used cells from mice carrying the Cd23Cre transgene, loxP-flanked Bax alleles and a germline deletion of Bak1 ^46^. Mature B cells, expressing CD23, will lack both BAX and BAK proteins, rendering them incapable of activating the distal stages of apoptosis and are referred to as Bax/Bak KO ^46, 47, 48, 49^. B cells from Bak1-/- mice can undergo normal apoptosis (Supplementary Fig. 2a) due to the functional redundancy of BAX and BAK and were therefore used as the control for all Bax/Bak KO studies. Bax/Bak KO mice develop relatively normal mature follicular B cells that, due to impaired BAFF competition are typically, on average, smaller than wildtype ^46^. Here we use density gradients to purify cells of similar size, and as such no differences were observed between control and Bax/Bak KO cell size (Supplementary Fig. 2b). In response to CpG stimulation, Bak1-/- control and Bax/Bak KO B cell numbers were initially equivalent, however after 72h, control cells were lost, while Bax/Bak KO numbers remained alive (Fig. 2a). These differences are solely explained by the extended survival of Bax/Bak KO B cells (Fig. 2b), as mean division number (MDN) was comparable to control B cells (Fig. 2c). Due to the inability of Bax/Bak KO B cells to undergo apoptosis, the small proportion of cells not activated after exposure to CpG, that die in control cultures, persist in division zero thus skewing the MDN. Quantifying the MDN of dividing cells, excluding division zero, revealed identical proliferation of control and Bax/Bak KO B cells (Fig. 2d), indicating equivalent activation, and therefore validating the Bax/Bak KO system as a useful and representative tool for the quantification of BCL-2 family protein expression in cells that would normally have died.

**Figure 2.**
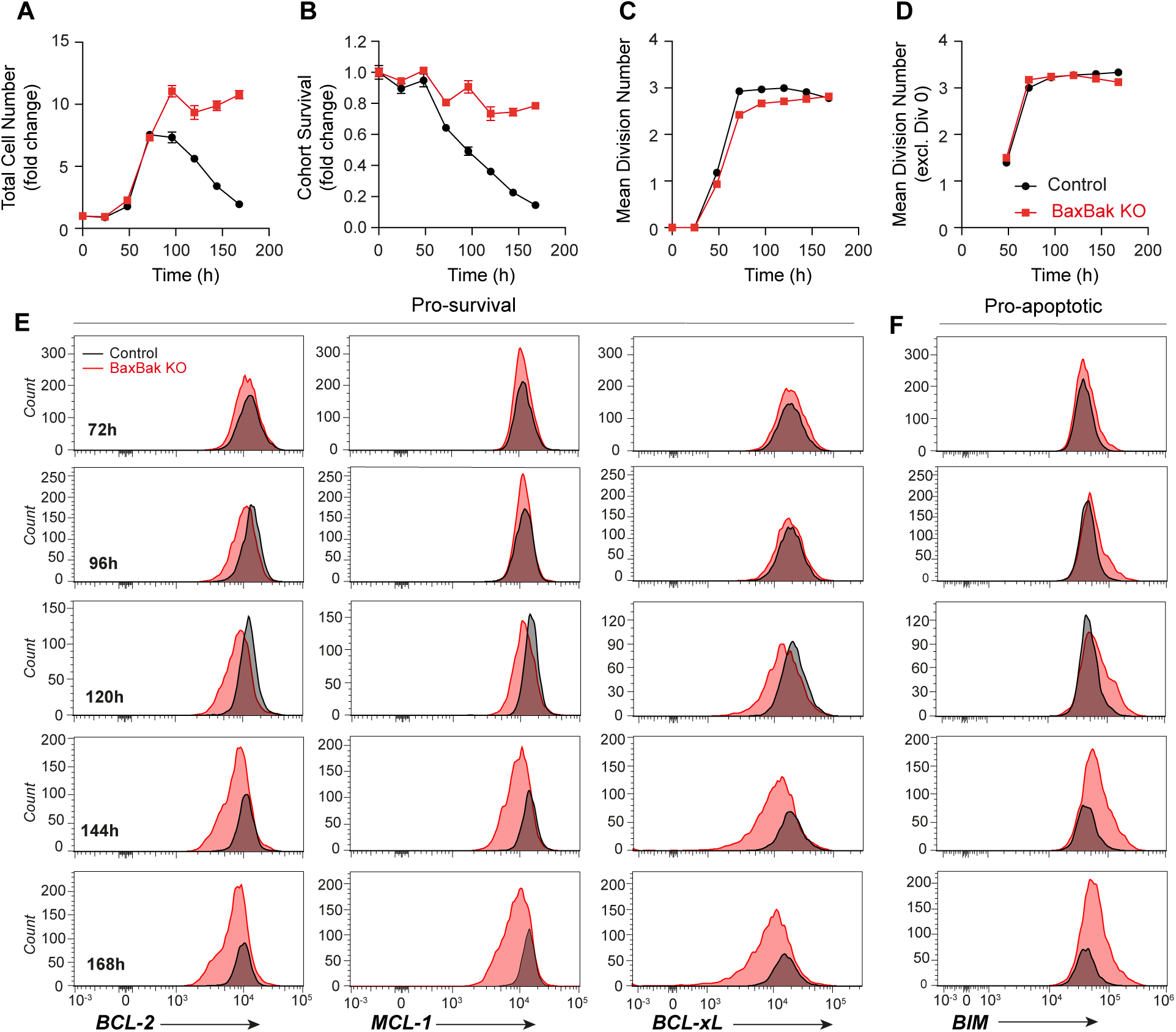
Blocking apoptosis reveals BCL-2 family protein censorship over time. Control and Bax/Bak KO B cell responses to CpG stimulation including (**a**) total cell number, (**b**) cohort survival, (**c**) mean division number and (**d**) mean division number excluding undivided cells were analyzed over time. (**e-f**) Flow cytometry analyses of BCL-2 family protein expression after CpG activation in control and Bax/Bak KO B cells. (**e**) Overlaid expression histograms of pro-survival proteins BCL-2, MCL-1 and BCL-xL and (**f**) pro-apoptotic protein BIM over time. Data in a-d are represented as mean ± SEM, and e-f are representative plots of triplicate samples. Data representative of three or more independent experiments.

To explore the molecular mechanics of timed cell death we measured the levels of key BCL-2 family proteins over time by flow cytometry. In addition to assessing the distributions of BCL-2, MCL-1, BCL-xL and BIM expression in the population during the death window (Fig. 2e-f) the population geometric mean fluorescence intensity (gMFI) over time was assessed from time of activation (Supplementary Fig. 2b). Following activation, pro-survival proteins BCL-2, MCL-1 and BCL-xL followed similar patterns of expression, with gMFI peaking at 48 hours and reducing over time, with reduced expression of all pro-survival proteins observed in Bax/Bak KO B cells compared to control cells after 72 hours. In contrast, the expression of pro-apoptotic protein BIM was consistently higher in Bax/Bak KO B cells during this time (Supplementary Fig. 2b).

Therefore, in the absence of cell death, pro-survival proteins gradually decreased over time and the pro-apoptotic protein, BIM, gradually increased, arguing against sudden loss, or rapid expression, of one or more proteins as a trigger for apoptosis. Further, as cells were CTV labelled, we could track the pattern of rise and fall in cells from different generations (Supplementary Fig. 2d). This indicated that division was not altering the changing pattern of expression. Thus, the kinetic control over expression appeared heritable, consistent with the clonal concordance in cell death ^5^.

The overlaid expression histograms demonstrate a time-dependent accumulation of Bax/Bak KO B cells with low levels of pro-survival proteins (Fig. 2e) and elevated levels of BIM (Fig. 2f). These data indicated that cells with low levels of pro-survival protein expression and/or high levels of BIM are lost from the wild type cultures via apoptosis. Figure 2 also shows that, with time, the levels of pro-survival molecules BCL-2, MCL-1 and BCL-xL are diminishing while pro-apoptotic BIM expression increases, thus tipping the balance in favor of apoptosis. Importantly, the upper expression boundaries for the pro-survival proteins and the lower limits of BIM expression were consistent between control and Bax/Bak KO B cells. These results are consistent with the hypothesis that a critical threshold of combined BCL-2 family protein expression is required to preserve B cell survival. By this hypothesis, kinetic changes in protein levels during the death phase means cells will exhibit a distribution of death times within the population, determined by the combined expression of individual proteins in each cell in relation to the triggering threshold. We sought to test this prediction directly by modelling the fate of each individual cell.

### A predictive cell death model incorporating an ensemble of BCL-2 family proteins relative to a critical survival threshold

We developed a quantitative model where the combined amounts of BCL-2 family proteins could predict cell survival outcomes, explaining and recreating the censored protein levels observed in surviving control cells. We refer to such a protein combination as an ‘ensemble’, and a model whereby the ensemble must remain above a threshold for cell survival as the Ensemble-Threshold (ET) model. The precise mathematical definition of this protein ensemble will be explored below. Figure 3a provides an overview of the model and how it was fitted to our experimental data. Details of the fitting workflow are provided in the methods.

**Figure 3.**
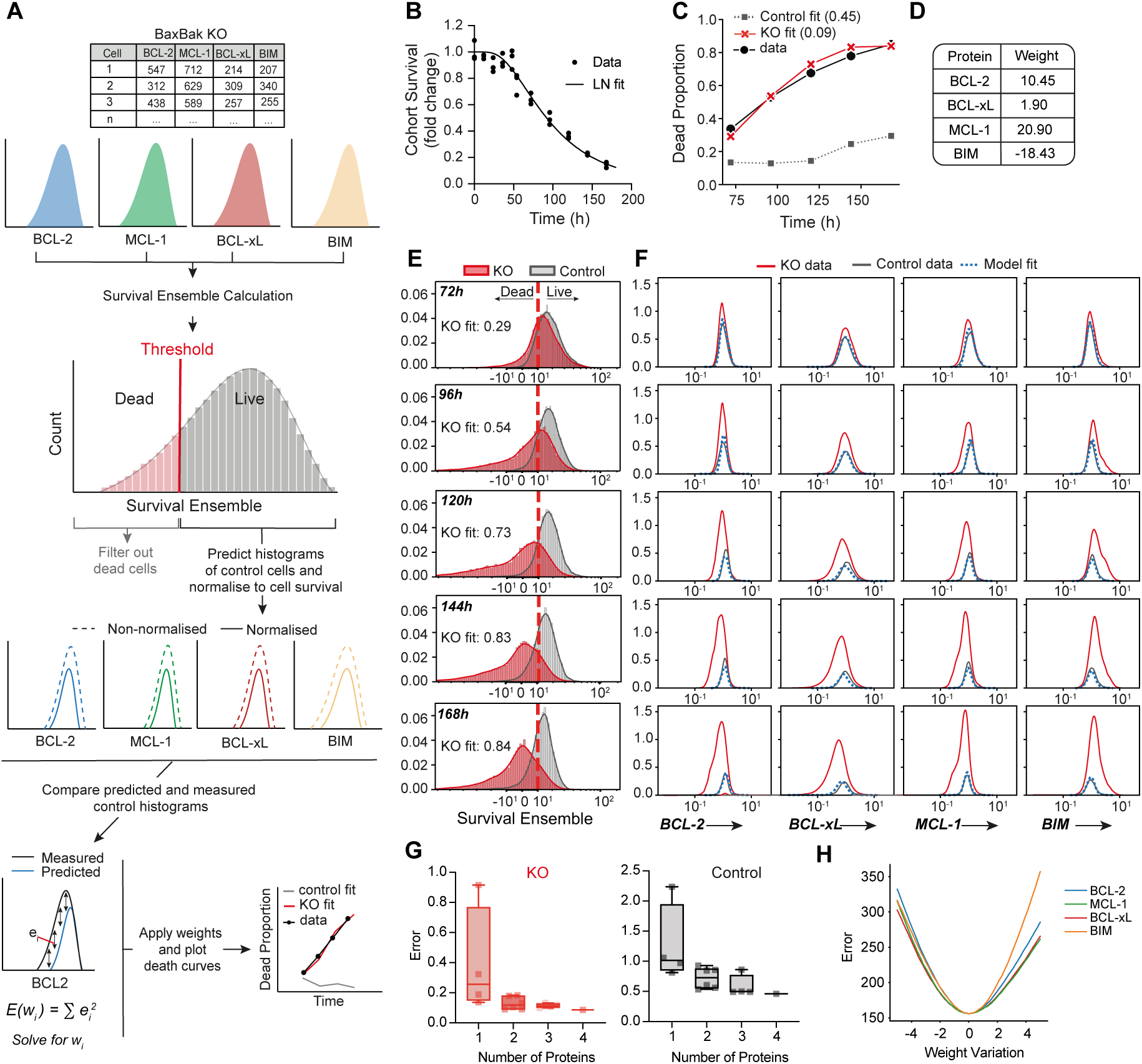
The Ensemble-Threshold Model: survival ensemble loss below a threshold recapitulates BCL-2 family protein expression and death curves. **(a)** Schematic of the Ensemble-Threshold model and fitting algorithm. (b-e) Ensemble-Threshold (SUM) model analysis of CpG activated control and Bax/Bak KO B cell survival and flow cytometry expression data, as shown in Fig 2. (**b**) Lognormal fits of control cohort survival data were performed to determine measured cell death over time. (**c**) Cell death predictions were calculated by applying (**d**) fitted protein weights to measured BCL-2 family expression levels and (**e**) survival ensemble expression over time was quantified. As a measure of fitting, survival curve error is included in the legend of cell death prediction plots. (**f**) The ET model fits of BCL-2 family protein expression overlaid with observed data. (**g**) The Bax/Bak KO and control cell survival curve errors when varying the number of proteins incorporated into ET model analysis. (**h**) The sensitivity of fitting to variation of calculated protein weights. Data representative of three independent experiments.

In principle, the mathematical function describing the survival ensemble value could be complex; however, to keep unknown parameters to a minimum and in the absence of other information, we initially used a simple arithmetic approach – a weighted linear sum (Eq. 2). The reasoning behind this approach was the known role of pro-survival proteins to ‘buffer’ the ability of BH3-only proteins, such as BIM, to bring about BAX/BAK activation and mitochondrial outer membrane permeabilization (MOMP) ^18, 20, 22, 50^. As BIM is capable of inhibiting all of the pro-survival molecules tested here ^20^, we assume a positive weight for pro-survival BCL-2 family proteins, is balanced against a negative weight for pro-apoptotic BIM expression.

To test the linear sum ET model, survival, and BCL-2 protein family expression data of CpG stimulated control and Bax/Bak KO B cells were analyzed. Lognormal death curves were directly determined by fitting to precursor cohort survival data (Fig. 3b). Fitting the linear sum ET model resulted in Bax/Bak KO death curve fits that were almost identical to the observed control cell death over time (Fig. 3c). Moreover, control B cell fits remained close to zero suggested the ET model is correct (Fig. 3c). The protein weights that provided the best fit (Fig. 3d) were applied to the measured BCL-2 family expression to determine and visualize survival ensembles over time (Fig. 3e). The Bax/Bak KO survival ensemble distributions shifted with time, with an increasing proportion of cells falling below the survival threshold over time. Control survival ensembles reduced slightly with time; however, they were clearly bounded by the threshold, with the vast majority above the survival threshold at each timepoint. The ET model prediction of control B cell protein expression histograms very closely mirrored the measured data for the various BCL-2 family members over time and replicated the censorship patterns observed compared to the measured Bax/Bak KO protein expression (Fig. 3f). Together these results indicate that Bax/Bak KO B cells replicate the conditions for triggering apoptosis operating in control cells and that information provided by just four of the more than 15 potentially expressed BCL-2 family members is sufficient to explain the broadly distributed pattern of cell loss over time. Thus, an ensemble based on a simple weighted sum provides a surprisingly accurate model that recapitulates observed censored protein expression and B cell death curves.

We explored the importance of each protein to the ET model fits by varying the number and combination of proteins included in the ensemble. We calculated the error in the control and Bax/Bak KO death curves for each of 15 possible protein combinations (Fig. 3g). The most accurate death curve fits were found when incorporating the expression data of all proteins measured, with increased error observed as the number of proteins included was reduced. The sensitivity of fitting to changes in level of optimal protein weights, again indicate that each of the four proteins are contributing to the best solution and the weight values, once determined, are well constrained (Fig. 3h).

The ability of the ET model to fit multiple data sets with a single set of protein weights was investigated. The simultaneous ET model fitting resulted in appropriate fits across three data sets, with accurate predictions of death curves (Supplementary Fig. 3a). The protein weights (Supplementary Fig. 3b), once determined, were again well constrained (Supplementary Fig. 3c). Survival ensemble and BCL-2 family expression predictions were also accurate (Supplementary Fig. 3d-f). These results suggest a universal solution may apply if experimental noise can be minimized and absolute levels of cell proteins determined.

We explored alternative functions for determining the survival ensemble value. A ratio of pro-survival to pro-apoptotic BCL-2 protein expression (Eq 3) performed well (Supplementary Fig. 4) but the linear sum model consistently produced more accurate fits across multiple independent experiments. Together, the level of accuracy of death curve fits suggests the linear sum method reflects the underlying biological mechanism well and clearly demonstrates that the timed reduction in BCL-2 family ensemble expression below a critical threshold is regulating the death times of CpG activated B cells.

### The ET model predicts the effect of BH3 mimetics on B cell survival and protein

How might the ET model accommodate the ability of BH3 mimetics to accelerate times to die (Fig. 1)? We hypothesized that BH3 mimetics alter the effective weight of the targeted protein and therefore lower the ensemble values at each time point. This drug-induced shift in the ensemble values will increase the proportion of cells that fall below the threshold. Given the interest in predicting the efficacy of these drugs clinically, we examined this hypothesis as a further test of our ET model.

To generate data for modelling, cohort survival and BCL-2 family protein expression was measured for control and Bax/Bak KO B cells following CpG stimulation and subsequent MCL-1 inhibition. As expected, MCL-1 inhibition resulted in a dose-dependent reduction of B cell survival of control, but not Bax/Bak KO, B cells (Fig. 4a). Despite a reported effect of MCL1 protein stabilization (47), BH3 mimetic treatment did not alter the expression of target molecules in Bax/Bak KO cells (Supplementary Fig. 5a), indicating an impact on protein function, not stability in this case. A key feature of the ET model was the accrual of Bax/Bak KO B cells with low levels of pro-survival proteins and high pro-apoptotic protein expression over time. Accordingly, the undrugged CpG stimulated B cells displayed the typical skewed protein expression patterns and, importantly, this feature was exaggerated following MCL-1 inhibition (Fig. 4b). These findings indicate that MCL-1i hastens cell death kinetics via a functional inhibition of MCL-1, exacerbating the censorship of control cells at the lower end of the expression distribution.

**Figure 4.**
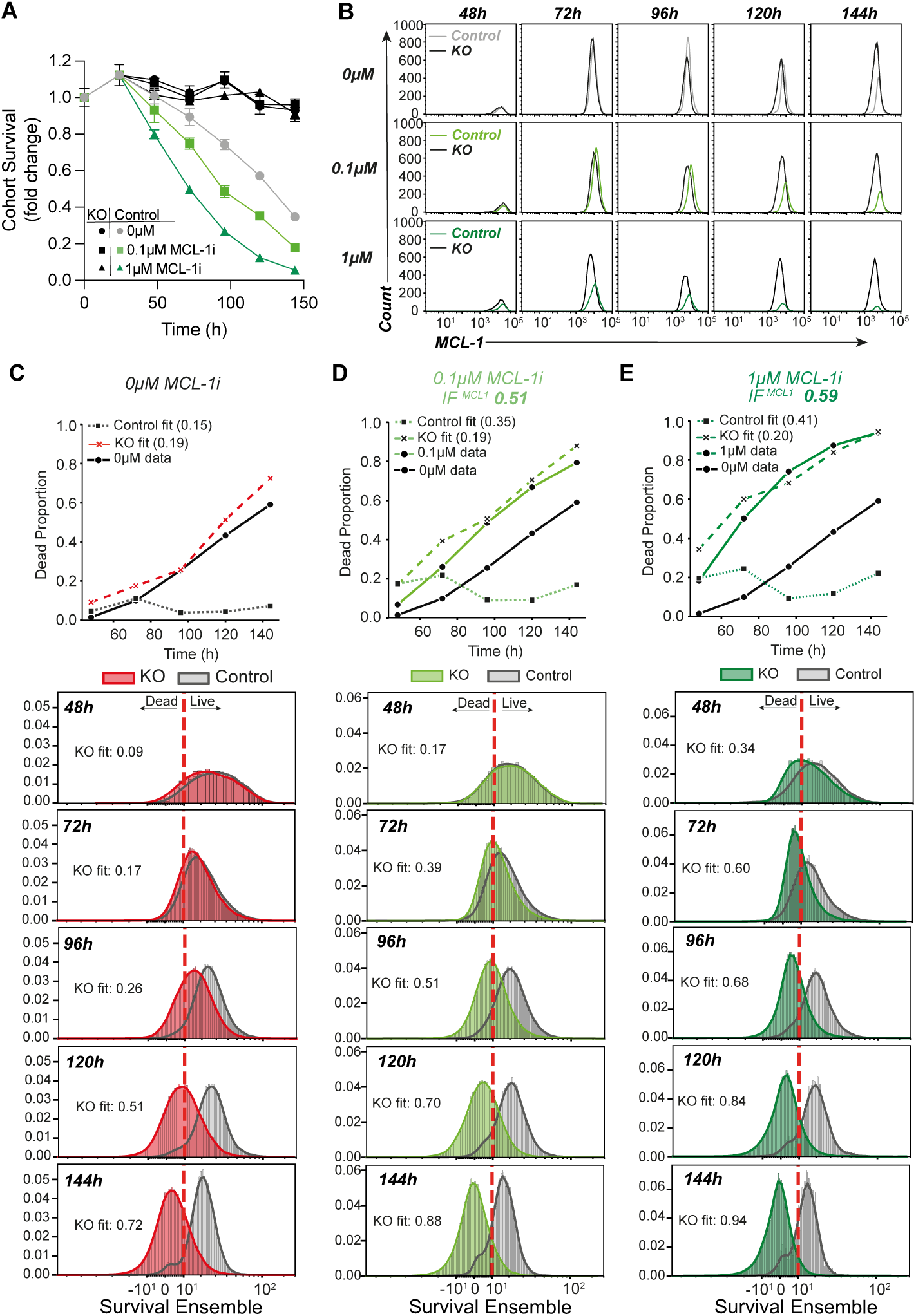
ET model explains BH3 mimetic effects on survival by reducing weight of target protein. **(a)** Cohort survival of control and Bax/Bak KO B cells following CpG activation and treatment with MCL-1i from 24h. Data represented as mean ± SEM. (**b**) MCL-1 protein expression was assessed in Bax/Bak KO cells following MCL-1i treatment. Data are representative plots from triplicate samples. (**c**) ET model analysis of undrugged CpG stimulated control and Bax/Bak KO B cells. (**d-e**) ET model death curve fits, survival ensemble expression over time and calculated IF values were quantified following (**d**) 0.1mM MCL-1i or (**e**) 1mM MCL-1i treatment. As a measure of fitting, survival curve error is included in the legend of cell death prediction plots. Data are representative of two independent experiments.

To identify the inhibitory impact of BH3 mimetic treatment on survival ensembles, we introduced a new parameter to reduce the effective weight of the targeted protein, by a fraction (Eq 4). Varying this inhibition factor (IF) between 0 and 1 will enable the effective influence of the target protein on the survival ensemble to be titrated from zero to full activity, respectively. To test this simple model of drug action, we applied the protein weights determined by ET model fitting of the undrugged response (Fig. 4c, Supplementary Fig. 5b-c) and then fitted the remaining IF parameter to the drug responses. A dose-dependent effect on the IF applied to MCL-1 expression following MCL-1 inhibition was calculated, with 0.1μM and 1μM MCL-1i resulting in an IF of 0.51 and 0.59, respectively. Accurate ET model death curve predictions following MCL-1 inhibition were observed (Fig. 4d-e) along with accurate predictions of the censored control cell BCL-2 family expression (Supplementary Fig. 5d-e). Analogous results were obtained for ET model analysis of cells exposed to BCL-2 inhibition, with incorporation of dose dependent IFs able to recapitulate protein expression, survival ensemble loss and death curve fits following BCL-2 inhibition of CpG stimulated B cells (Supplementary Fig. 6a-f).

If correct, our ET model predicts that altered times to die induced by BH3 mimetics, will be the same, regardless of the time the drug is added. There should be no memory, or accumulation, of drug effects over time. This prediction was supported by experiments. Similar effects of BH3 mimetic treatment at various times post CpG activation were observed, with immediate loss of a proportion of cells followed by gradual loss over time as cells followed their new death time (Supplementary Fig. 6g). Therefore, we conclude that the effect of BH3 mimetics to reduce the functional survival ensemble and alter B cell death times is independent of when the protein inhibition occurs post activation.

### The ET model predicts death in other cell systems

Having described a protein ensemble-threshold mechanism for activated B cell death time regulation, we asked whether related mechanisms could explain the varied death times of other cell systems. We applied the model to data from unstimulated B cells (Fig. 5a, b) and CD4+ T cell following IL-2 withdrawal (Fig. 5c, d).

**Figure 5.**
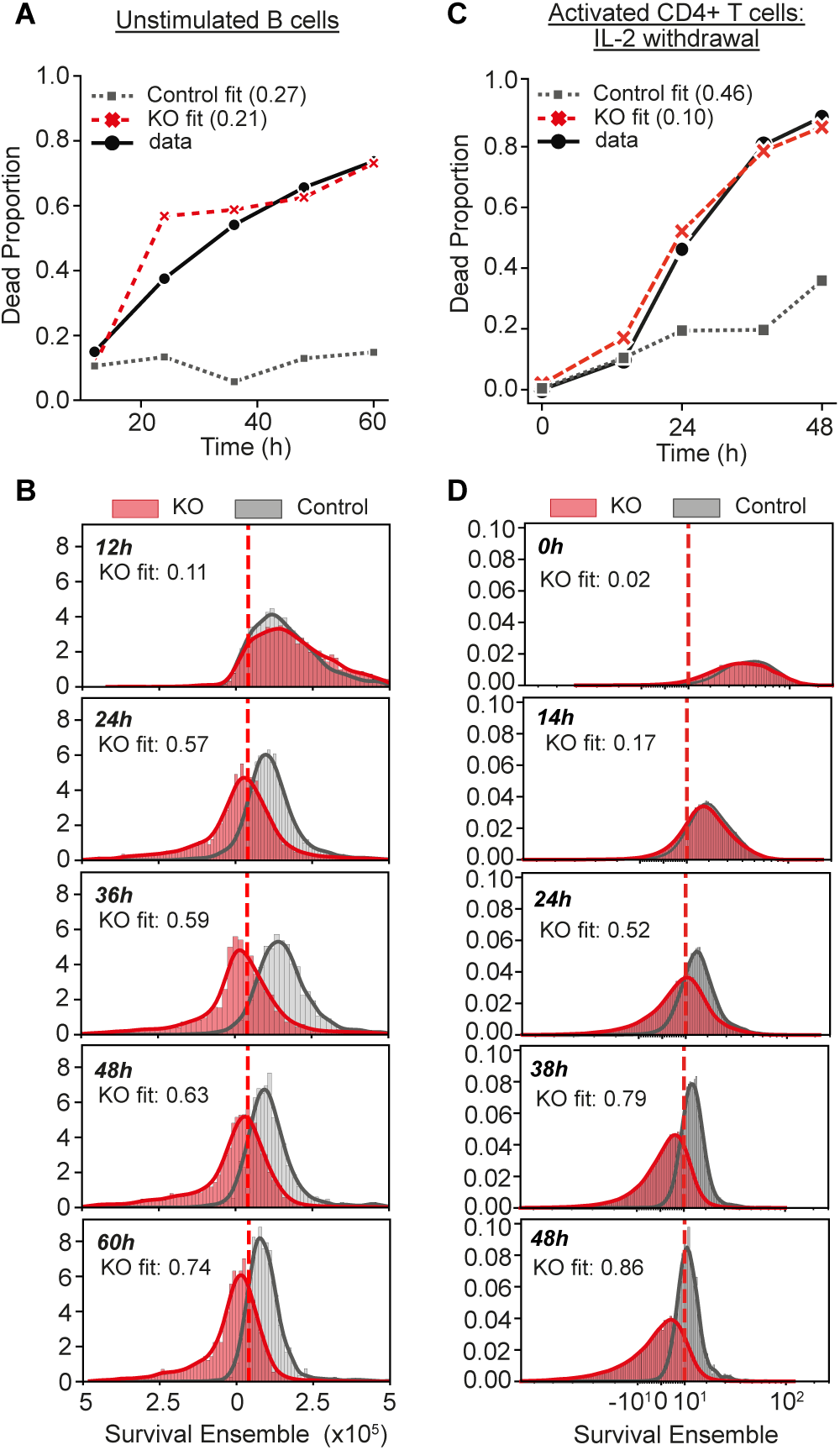
ET cell death mechanism applied across cell types and activation state. ET model analysis of (**a-b**) control and Bax/Bak KO unstimulated B cells over time and (**c-d**) control and Bax/Bak KO activated CD4^+^T cells following IL-2 withdrawal including (**a, c**) cell death curves and (**b, d**) survival ensemble expression over time. As a measure of fitting quality, survival curve error is included in the legend of cell death prediction plots. Data are representative of two independent experiments.

For B cells Bax/Bak KO and control cells were maintained in culture without activation. As expected, unstimulated Bax/Bak KO cells remained alive while WT B cells died rapidly over time with half-life of around 24 h (Supplementary Fig. 7a). Survival data and the expression of BCL-2 family proteins were measured and analyzed using the ET model. Cell death curve predictions from fitting Bax/Bak KO data replicated the observed death rate of control cells (Fig. 5a). The fitted protein weights (Supplementary Fig. 7b) were applied to the measured expression values to determine and visualize survival ensemble expression changes over time (Fig. 5b). The proportion of unstimulated Bax/Bak KO B cells whose survival ensembles were below the survival threshold increased over time, accurately reflecting the observed death of unstimulated control cells while predicting the BCL-2 family protein expression over time (Supplementary Fig. 7c). These results demonstrate the protein ensemble - threshold mechanics described for activated B cell death also apply to regulation of unstimulated B cell death.

IL-2 transmits a potent T cell survival promoting signal to activated cells and its removal leads to rapid cell death ^42, 51^. CD4Cre Bak1-/- Bax fl/fl (Bax/Bak KO) and control (Bak1-/-) CD4+ T cells were stimulated with aCD3/aCD28 and IL-2 for 48h, after which cells were maintained in 100U/ml IL-2. At 96h after initial activation, Bax/Bak KO and control CD4+ T cells were split into fresh media with or without IL-2. Activation of control and Bax/Bak KO CD4+ T cells induced substantial expansion (Supplementary Fig. 7d). Both control and Bax/Bak KO CD4+ T cells continued to proliferate in response to IL-2 supplementation. Following IL-2 withdrawal, Bax/Bak KO CD4+ T cell numbers plateaued, reflecting a cessation of division, whereas control CD4+ T cells numbers markedly decreased over time indicating, in addition to arrested division, cells died with half-life around 24 h. The expression of BCL-2 family proteins was comparable in control and Bax/Bak KO CD4+ T cells at the time of IL-2 withdrawal (Supplementary Fig 7e). As CD4+ T cell proliferation was beyond the resolution of CTV propidium iodide (PI) was used to determine cell viability from the point of IL-2 withdrawal, showing that control CD4+ T cells rapidly died whereas Bax/Bak KO CD4+ T cells maintained survival (Supplementary Fig. 7f).

The CD4+ T cell survival data and the expression of BCL-2 family proteins over time after IL-2 withdrawal was analyzed using the ET model. Cell death curve predictions from fitting Bax/Bak KO data replicated the observed death rate of control cells (Fig. 5c). The fitted protein weights (Supplementary Fig. 7g) were applied to the measured expression values to determine and visualize survival ensemble expression changes over time (Fig. 5d). The proportion of Bax/Bak KO CD4+ T cells whose survival ensembles were below the threshold increased over time, accurately reflecting the observed death of control cells. Importantly, control ensembles remained above or around the set survival threshold, with those below being lost from the analysis. The ET model also accurately predicted the BCL-2 family protein expression of control CD4+ T cells over time (Supplementary Fig. 7h). Together, these results demonstrate the ET model and the underlying protein ensemble - threshold mechanics described for B cells are appropriate to explain the loss of CD4+ T cells in response to IL-2 withdrawal.

## DISCUSSION

Understanding how lymphocytes integrate and process signals to control cellular fates appropriately and effectively is a subject of intense interest and is fundamental to immunology. Increasingly, it is clear that these processes are mathematically precise, and a quantitative understanding of immune cell response dynamics can be used to provide insight into the underlying molecular processes driving immune responses ^4, 5, 7, 11, 12, 13^. The major goal of this study was to develop and apply quantitative methods to identify the molecular machinery controlling clonally heritable death times that are a striking feature of long-term filming studies of B and T cells ^5, 10^. We explored the central hypothesis that the heritable death timer is regulated by the expression of critical molecules reaching, or falling below, threshold levels required for survival.

As the role of BCL-2 family proteins in cell survival is well known, we set out to relate the levels of these proteins to the time of initiation of apoptosis by activated B cells. Thus, we sought a quantitative implementation of the concept of apoptotic priming, where cells can vary in their apparent ‘distance’ from the apoptotic threshold^34, 35, 36^. In our quantitative system, numerous features are identifiable. Naïve B cells, placed in culture die rapidly ^4^. CpG activation reprograms their survival and extends the half-life to around 96h. B cell death times are variable and altered by BH3 mimetic treatment and genetic ablation of BIM, demonstrating the quantitative nature of the death time mechanism. Notably, the timed survival mechanism could be overridden by simultaneously inhibiting the function of BCL-2, MCL-1, BCL-xL identifying these three pro-survival proteins as the principal and essential drivers of the timing process and validating their use in further model building.

Our ability to investigate BCL-2 family expression over time in apoptosis-deficient B cells proved critical. The disparity in expression of BCL-2 family proteins between the apoptotic deficient Bax/Bak KO B cells and surviving control cells over time supported the hypothesis that a critical threshold of combined ensemble expression of BCL-2 family proteins was required for continued cell survival. As such, the default loss of expression over time triggered death if the ensemble breached a threshold. Further, the broad variation in expression of each protein meant each cell was on a unique path with its own time to die that resulted in surviving cells presenting striking censored protein features. These patterns of expression change were not altered by division explaining the remarkable symmetry in familial clonal fate^5^. We sought rules of BCL-2 family regulation and addition to generate a quantitative model of B cell death. We found an effective protein ensemble function for the combination of BCL-2 family expression that recapitulated observed protein expression patterns in surviving cells and correctly accounted for timed death curves. The cooperative action of pro- and anti-apoptotic proteins to regulate time to die extends and complements earlier studies ^25, 33, 52^. The most accurate and consistent estimates of cell death in the ET model were obtained using a simple linear sum function, subtracting weighted BIM expression from the positively weighted sum of pro-survival molecule expression. This function reflects the action of pro-survival molecules to neutralize the activation of Bax and Bak and incorporates BIM in a manner consistent with both the direct ^22, 50, 53^ and indirect models ^18, 54^ of apoptosis. Encouragingly, ET model analysis of CD4 T cell survival following IL-2 withdrawal was sufficient to replicate the changes in BCL-2 family protein expression and accurately recapitulate observed cell death implying a general mechanism for lymphocytes.

Our method of ensemble calculation, based on the four limiting BCL-2 family proteins proved sufficient for robust and precise estimations of activated cell death in B and T cells. However, we anticipate prediction accuracy might be improved with additional components. In other cell systems measurements of further BCL-2 family proteins ^16, 55, 56, 57^ may be required to achieve a similar degree of accuracy as seen here for T and B cells. Further our model of lymphocyte control requires active survival maintenance by signals. When B and T cells are isolated, the signal deficient state leads to death over time. This default pattern of death, in our experimental model is principally due to the loss of expression of pro-survival proteins, and, to a lesser extent, the rise in pro-apoptotic protein BIM. These assumptions of the ET model can be extended to other cell models where appropriate. Homologues for key components of the cell death pathways, described here to regulate death times, are found in a wide range of organisms ^58, 59^. Thus, cell death is regulated by evolutionarily conserved control mechanisms, and we anticipate the mathematically precise rules identified here for B and CD4+ T cells may be applicable for additional cell types within other systems where the default is cell death unless survival signals are received.

We have been able to extend the ET model to reproduce effects of BH3 mimetics with a single additional parameter. The integration of a scalable, protein inhibition factor to the ET model was sufficient to provide accurate predictions of enhanced cell death and the resulting alterations to the censored protein expression levels. These mechanisms held true for combination BH3 mimetic treatment and when protein inhibition was introduced at different times post activation, thus supporting key predictions of the ET model. Owing to the ability of cells to rely on more than one BCL-2 family protein for survival, the combined use of multiple BH3 mimetic treatments is a thriving area of cancer research ^60, 61^. Where BCL-2 family expression can be measured, the ET model could be applied to predict the effects of combinations of BH3 mimetic treatment on tumor cells, providing critical information and potentially enhancing and personalizing the effective use of these drugs clinically.

Our results clearly indicate the fate of lymphocytes is strongly determined by the molecular levels of regulatory proteins, and importantly, variation of protein expression between cells ensures non-identical outcomes of cell fate within a population. Real-time regulation of critical protein levels by integrating signals, coupled to default changes to expression once signals wane, or are lost, explain the timer-like features we observe. We anticipate that multiple signals can be processed to add to the time for division, by expression of Myc, and the time for death, by expression of protein ensemble components, to alter the overall immune response dynamics. Combination signaling through BCR, TCR, cytokines and other costimuli will enhance the initial production rate of the critical proteins and alter the final dynamics in B and T cells, in a manner we expect to have been optimized during evolution to generate and match an appropriate rate of expansion and timed loss of cells to the level of threat ^62^. By this logic it is unsurprising that the molecular regulators found to control the cell death timer and division destiny timer are well known, evolutionarily conserved proteins, whose regulation is tightly controlled and frequently dysregulated in lymphoma.

Overall, this study builds on the larger goal of developing models of lymphocyte control that bridge molecular mechanisms and population dynamics ^4^ ^11^ ^10^ ^38^. Our results support the hypothesis that each independent cell fate module functions as a timer, controlled by the levels of key molecules. The model for cell death prediction developed here incorporates the mechanistic and stochastic features of cellular control into a rule set that can potentially recreate the population outcomes under complex signaling conditions. For effective models we can reduce complexity, focusing only on the key elements that govern the dynamics of the model. Here we established the BCL-2 family ensemble as the limiting element in death time regulation, that can be summarized in a manner that informs the operation of the death timer. Importantly, these discoveries provide a framework that we can now use to interrogate these mechanisms in other settings. In disorders such as primary immunodeficiency, autoimmunity, and cancer, characterizing the parameters contributing to dysregulated responses could indicate which molecular components are likely to be affected, thus, potentially informing therapeutic strategies.

## MATERIALS AND METHODS

### Mice

C57BL/6, BCL-2l11^-/-^ (Bim KO), Bak1^-/-^ (Bak KO) and Cd23^Cre^ Bak1^-/-^ Bax^fl/fl^ or Cd4^Cre^ Bak1^-/-^ Bax^fl/fl^ (Bax/Bak KO) mouse strains were maintained under specific pathogen-free conditions in the Walter and Eliza Hall Institute animal facility. Male and female mice were used between 6-14 weeks of age. Sex-matched, littermates or age matched controls were used. All procedures were performed as approved by the WEHI animal ethics committee. Mice carrying the Cd23^Cre^ Tg ^63^, the germline deletion of BCL-2l11 ^42^, Bak1 ^64^, the floxed alleles of Bax ^65^ and the combined Cd23^Cre^ Bak1^-/-^ Bak^fl/fl^ ^46^ have been previously described.

### B and T cell culture

B cells and CD4^+^ T cells were cultured in RPMI-1640 medium, supplemented with 10% (vol/vol) FBS, 10mM HEPES, 100U/ml penicillin, 100mg/ml streptomycin, 2mM Glutamax, 0.1mM non-essential amino acids, 1mM sodium pyruvate (all Invitrogen), and 50mM β-mercaptoethanol (Sigma). For division tracking, cells were labelled with 5mM CellTrace Violet (CTV, Invitrogen) in sterile PBS containing 0.1% Bovine Serum Albumin (PBS 0.1% BSA), according to manufacturer instructions.

Small resting B cells were isolated from spleen with a discontinuous Percoll gradient and subsequent purification using EasySep Mouse B cell Isolation Kit (StemCell Technologies). B cells were stimulated with 3mM CpG (Sigma), in 96 well flat-bottom plates at 10^4^ cells per well.

CD4^+^ T cells were isolated from peripheral lymph nodes (inguinal, axillary, brachial and superficial cervical) by negative selection using the EasySep Mouse CD4^+^ T cell Isolation Kit (StemCell Technologies). CD4^+^ T cells were stimulated in 96 well flat-bottomed plates coated with 20mg/ml αCD3 (clone 145-2C11, WEHI antibody facility), in the presence of 2mg/ml aCD28 (clone 37.51, WEHI antibody facility) and 100U/ml recombinant human IL-2 (Peprotech), in 96 well flat-bottom plates at 10^4^ cells per well. 48h post activation cells were harvested, washed, and replated in the presence of 100U/ml IL-2. 96h post initial activation T cell cultures were harvested, washed, and replated either with or without 100U/ml of fresh recombinant IL-2 and survival was monitored over time.

### Quantitative cell analysis

A known number of beads were added to samples immediately before cell counting analysis was performed on a FACSCanto II (BD Biosciences). Propidium iodide (0.2mg/ml, Sigma) was used for dead cell exclusion and the ratio of beads to live cells was used to determine the cell number in each sample.

Cohort survival and mean division number (MDN) were determined using a modification of the precursor cohort method previously described^3, 12, 13, 40^. The precursor cohort method removes the effect of proliferation on total cell numbers, thus providing an estimate of loss of original founder cells whose clonal family contributes to the overall response at a particular time. The total cohort number over time is indicative of the net survival features of the population. The MDN describes the average number of divisions the original cell cohort have completed.

To determine death time distributions, a one minus cumulative lognormal distribution function was used to fit experimental cohort survival data. The mean and variation parameters of the underlying lognormal distribution were iterated to find fits to measured survival with the smallest residual sum of squares. Where death time distributions were represented by the calculated lognormal curves and the corresponding one minus cumulative distribution replicated measured survival curves.

### Intracellular Antibody Staining

Cells were harvested at timepoints indicated and following incubation with fixable viability dye (Live/Dead Fixable Yellow or Fixable Viability Dye eFluor780) in PBS (1:1000, Thermo Fisher Scientific), were washed and resuspended in fixation buffer (0.5% paraformaldehyde, 0.2% Tween-20 and 0.1% bovine serum albumin in PBS) for 24h. Cells were washed and resuspended in FACS buffer (PBS with 1%BSA, 1mM EDTA and 0.05% sodium azide) and stored at 4°C until all timepoints were harvested. Cells were stained at 4°C for 45 minutes with the following antibodies: anti-BIM FITC (1:400, clone C35, WEHI antibody facility); anti-Active Caspase 3 BV650 (1:200, clone C92-605, BD Biosciences); anti-BCL-xL PE (1:200, Clone E18, Abcam); anti-BCL-2 PE-Cy7(1:200, clone 10C4, BD Biosciences) and anti-MCL-1 A647 (1:1000, clone 19C4-15, WEHI antibody facility) and analysed on a Fortessa X20 (BD Biosciences) or Cytek Aurora spectral flow cytometer (Cytek Biosciences). Data was acquired and analysed using SpectroFlo and FlowJo software.

### Mathematical Modeling

#### Cyton Modeling

For analysis of B cell responses, we fitted a version of the Cyton2 model parameterised as described in Cheon et al ^11^. This model describes lymphocyte population dynamics after stimulation as resulting from the independent regulation of three competing cellular timers. The timers are parameterised by three random variables: (i) the time to first division; (ii) the division destiny time, and (iii) the time to cell death and the constant for the time between subsequent divisions. Random variables are drawn from lognormal distributions which are determined as described in Cheon et al ^11^. For each time-course data set, we aimed to determine the best-fit lognormal distributions for times to first division, division destiny time and death times.

### Ensemble-Threshold Model

For each cell, expression of BCL-2 family proteins was determined by intracellular staining and flow cytometric analysis. Exclusion of dead and apoptotic cells was performed using fixable viability dyes and staining for active caspase 3. Protein levels for each experiment were normalized by the geometric mean of the Bax/Bak KO protein level at 72 hours, the beginning of the death phase and where protein expression was comparable for control and Bax/Bak KO cells.

The Ensemble-Threshold model is defined in terms of an ensemble function, e(p_i_; w_i_), which has weights, wi, and takes protein level measurements, pi, as input and i represents the proteins BCL-2, BCL-xL, MCL-1 and BIM. A cell is presumed to be dead if

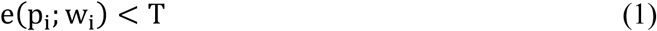

for some threshold T.

We have examined two simple and parsimonious functions for the ensemble function.

The first ensemble function we refer to as the linear sum model.

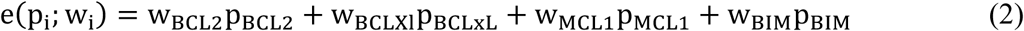

The function is expanded here explicitly in terms of the proteins. The weights for the pro-survival proteins are constrained to be positive and the weight for BIM is constrained to be negative. For this function, the weights and threshold can be uniformly rescaled without changing the live/dead determination for a cell, therefore the threshold was fixed at 10, without loss of generality.

The second ensemble function we examined was a ratio model.

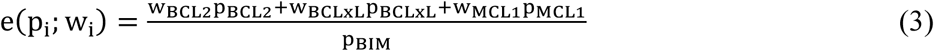

In this case, there are only three independent weights and, as before, the threshold was fixed at 10.

As is usual when fitting model parameters to experimental data, we created an error function that measures the difference between the model prediction and the data. For each cell measurement in the Bax/Bak KO data set, we calculated the ensemble value according to the particular ensemble function being tested (Eq 2 or 3). Bax/Bak KO cells below the threshold were presumed dead and were filtered out. New predicted protein histograms were constructed from the cells predicted to be alive. To make a quantitative comparison between control and Bax/Bak KO histograms they were normalized to relative cell numbers. We fitted a lognormal cumulative distribution function to the control cohort survival curves to calculate relative cell survival and used this to normalize the area under control protein level histograms. The area under the input Bax/Bak KO histograms were normalized to one. An error for each histogram pair was calculated by summing the square of the difference in the size of each histogram bin. This error was calculated for each protein, at each time point, and these errors were summed to create the model error function. When fitting several experiments simultaneously (fig. S3G-L) the error was the sum of the errors from each experiment.

When modelling the effect of BH3 mimetic, Eq 2 is modified to include an additional inhibition factor, IFi for protein i. For example, when modelling the effect of the MCL-1 inhibitor, Eq 2 becomes

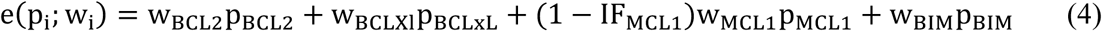

When fitting drug data, weights were first fitted to the undrugged control data and then fixed. The single remaining inhibition factor parameter was then fitted to the drugged data separately for each concentration.

Model fitting and plotting was done with Python 3.10 and errors were minimized using the differential evolution algorithm implemented in the Scipy Python library. This is a global optimization algorithm that takes parameter bounds as the primary input. The bounds for the pro-survival proteins were [0, B] and [-B, 0] for BIM (an exception is described below), where B was significantly larger than the eventual best parameter value. This algorithm has a stochastic component; however, parameter fits were reproducible within the specified tolerances.

Fig 3G was created by fitting linear sum ensemble models for all 15 protein combinations. In the case of the model with only BIM, a negative weight makes no sense so for these fits the BIM bound was [-B, B] for all combinations.

Ensemble values had a large dynamic range and were negative in some cases. For this reason, the abscissa of the ensemble histogram plots was formatted using the Logicle algorithm ^67^ implemented in the FlowCal 1.3.0 library ^68^. We have made fitting code publicly available (DOI: 10.5281/zenodo.18654246). The repository contains exported FACS data from the experiments presented here and scripts to reproduce all modelling figures.

## DATA AVAILABILITY

All data supporting the findings in this study are available within the main text, Supplementary, supplementary materials or contained within a publicly accessible GitHub repository (DOI: 10.5281/zenodo.18654246). All other data supporting the findings of this study are available from the corresponding authors on request

## CODE AVAILABILITY

All fitting code is made available, with data, scripts and instructions required to reproduce all modeling figures within the main text and extended material also available within the GitHub repository (DOI: 10.5281/zenodo.18654246).

## ACKNOWLEDGMENTS

We would like to thank all members of the Hodgkin and Gray laboratories for thoughtful discussions and suggestions; Stephane Chappaz and Benjamin Kile for the *CD23Cre Baxfl/fl Bak1-/- and CD4Cre Baxfl/fl Bak1-/-* mice; the Flow Cytometry Facility and Animal Facility at WEHI for technical assistance; Funding: This work was supported by the National Health and Medical Research Council of Australia (Project Grant 1164800 and Investigator Grant 1176588 to P.D.H.), Victorian State Government Operational Infrastructure Support, and Australian Government NHMRC Independent Research Institutes Infrastructure Support Scheme (361646). D.H.D.G. is supported by Australian NHMRC Fellowships/grant (1090236, 1158024 and 145888), The Medical Advances Without Animals Trust (MAWA) and Cancer Council Victoria grants-in-aid (1146518 and 1102104). C.E.T is supported by Victorian Cancer Agency Mid-Career (Fellowship: MCRF20026), NHMRC (2002618), Perpetual Impact Philanthropy (IPAP2019/1437) and Leukemia Foundation. M.R. was the recipient of a WEHI Internal Scholarship and a WEHI Edith Moffat Scholarship.

## AUTHOR CONTRIBUTIONS

M.R, S.H and P.D.H conceived and designed the study. M.R, S.H and P.D.H designed experiments, analyzed and interpreted experimental data. C.E.T and D.H.D.G developed staining protocols and interpreted experimental results. M.R and N.G performed the experiments. M.R, E.T, E.D, M.R.D and P.D.H analyzed data, tested, and constructed the mathematical models. M.R, E.T and P.D.H. prepared the figures and wrote the manuscript with input from all authors.

## REFERENCES

1. Iwasaki, A. & Medzhitov, R. Control of adaptive immunity by the innate immune system. Nature Immunology 16, 343–353 (2015).

2. Kaech, S.M. & Ahmed, R. Memory CD8+ T cell differentiation: initial antigen encounter triggers a developmental program in naïve cells. Nat Immunol 2, 415–422 (2001).

3. Gett, A.V. & Hodgkin, P.D. A cellular calculus for signal integration by T cells. Nat Immunol 1, 239–244 (2000).

4. Hawkins, E.D., Turner, M.L., Dowling, M.R., Gend, C.v. & Hodgkin, P.D. A model of immune regulation as a consequence of randomized lymphocyte division and death times. PNAS 104, 5032–5037 (2007).

5. Hawkins, E.D., Markham, J.F., McGuinness, L.P. & Hodgkin, P.D. A single-cell pedigree analysis of alternative stochastic lymphocyte fates. Proc. Natl. Acad. Sci. U. S. A. 106, 13457–13462 (2009).

6. Duffy, K.R. et al. Activation-induced B cell fates are selected by intracellular stochastic competition. Science (New York, N.Y.) 335, 338–341 (2012).

7. Marchingo, J.M. et al. T-cell stimuli independently sum to regulate an inherited clonal division fate. Nat. Commun. 7, 12 (2016).

8. Rush, J.S. & Hodgkin, P.D. B cells activated via CD40 and IL-4 undergo a division burst but require continued stimulation to maintain division, survival and differentiation. European Journal of Immunology 31, 1150–1159 (2001).

9. Turner, M.L., Hawkins, E.D. & Hodgkin, P.D. Quantitative regulation of B cell division destiny by signal strength. J. Immunol. 181, 374–382 (2008).

10. Mitchell, S., Roy, K., Zangle, T.A. & Hoffmann, A. Nongenetic origins of cell-to-cell variability in B lymphocyte proliferation. Proc Natl Acad Sci U S A 115, E2888–E2897 (2018).

11. Cheon, H. et al. Cyton2: A Model of Immune Cell Population Dynamics That Includes Familial Instructional Inheritance. Frontiers in Bioinformatics 1 (2021).

12. Heinzel, S. et al. A Myc-dependent division timer complements a cell-death timer to regulate T cell and B cell responses. Nat Immunol 18, 96–103 (2017).

13. Marchingo, J.M. et al. T cell signaling. Antigen affinity, costimulation, and cytokine inputs sum linearly to amplify T cell expansion. Science (New York, N.Y.) 346, 1123–1127 (2014).

14. Czabotar, P.E., Lessene, G., Strasser, A. & Adams, J.M. Control of apoptosis by the BCL-2 protein family: implications for physiology and therapy. Nature reviews. Molecular cell biology 15, 49–63 (2014).

15. Strasser, A., Cory, S. & Adams, J.M. Deciphering the rules of programmed cell death to improve therapy of cancer and other diseases. EMBO J 30, 3667–3683 (2011).

16. Youle, R.J. & Strasser, A. The BCL-2 protein family: opposing activities that mediate cell death. Nature reviews. Molecular cell biology 9, 47–59 (2008).

17. Fletcher, J.I. et al. Apoptosis is triggered when prosurvival Bcl-2 proteins cannot restrain Bax. Proceedings of the National Academy of Sciences 105, 18081–18087 (2008).

18. Willis, S.N. et al. Apoptosis initiated when BH3 ligands engage multiple Bcl-2 homologs, not Bax or Bak. Science (New York, N.Y.) 315, 856–859 (2007).

19. Oltvai, Z.N., Milliman, C.L. & Korsmeyer, S.J. Bcl-2 heterodimerizes in vivo with a conserved homolog, Bax, that accelerates programmed cell death. Cell 74, 609–619 (1993).

20. Chen, L. et al. Differential targeting of prosurvival Bcl-2 proteins by their BH3-only ligands allows complementary apoptotic function. Molecular cell 17, 393–403 (2005).

21. Davids, M.S. & Letai, A. Targeting the B-cell lymphoma/leukemia 2 family in cancer. Journal of clinical oncology : official journal of the American Society of Clinical Oncology 30, 3127–3135 (2012).

22. Kuwana, T. et al. BH3 domains of BH3-only proteins differentially regulate Bax-mediated mitochondrial membrane permeabilization both directly and indirectly. Molecular cell 17, 525–535 (2005).

23. Moldoveanu, T., Follis, A.V., Kriwacki, R.W. & Green, D.R. Many players in BCL-2 family affairs. Trends in biochemical sciences 39, 101–111 (2014).

24. Opferman, J.T. et al. Development and maintenance of B and T lymphocytes requires antiapoptotic MCL-1. Nature 426, 671–676 (2003).

25. Carrington, E. et al. Anti-apoptotic proteins BCL-2, MCL-1 and A1 summate collectively to maintain survival of immune cell populations both in vitro and in vivo. Cell Death Differ 24, 878–888 (2017).

26. Lecky, E. et al. Sequential apoptotic and multiplexed proteomic evaluation of single cancer cells. Science Advances 9, eadg4128 (2023).

27. Niepel, M., Spencer, S.L. & Sorger, P.K. Non-genetic cell-to-cell variability and the consequences for pharmacology. Current Opinion in Chemical Biology 13, 556–561 (2009).

28. Sorger, P.K. et al. Measuring and modeling life-death decisions in single cells. The FASEB Journal 26, 228.221–228.221 (2012).

29. Ameisen, J.C. On the origin, evolution, and nature of programmed cell death: a timeline of four billion years. Cell Death & Differentiation 9, 367–393 (2002).

30. Raff, M.C. Social controls on cell survival and cell death. Nature 356, 397–400 (1992).

31. Vaux, D.L. Toward an understanding of the molecular mechanisms of physiological cell death. Proc Natl Acad Sci U S A 90, 786–789 (1993).

32. Ellis, R.E., Yuan, J.Y. & Horvitz, H.R. Mechanisms and functions of cell death. Annu Rev Cell Biol 7, 663–698 (1991).

33. Mason, K.D. et al. Programmed anuclear cell death delimits platelet life span. Cell 128, 1173–1186 (2007).

34. King, L.E. et al. Apoptotic priming is defined by the dynamic exchange of Bcl-2 proteins between mitochondria and cytosol. Cell Death & Differentiation 29, 2262–2274 (2022).

35. Ryan, J.A., Brunelle, J.K. & Letai, A. Heightened mitochondrial priming is the basis for apoptotic hypersensitivity of CD4+ CD8+ thymocytes. Proceedings of the National Academy of Sciences 107, 12895–12900 (2010).

36. Singh, R., Letai, A. & Sarosiek, K. Regulation of apoptosis in health and disease: the balancing act of BCL-2 family proteins. Nature reviews. Molecular cell biology 20, 175–193 (2019).

37. Hawkins, E.D. et al. Quantal and graded stimulation of B lymphocytes as alternative strategies for regulating adaptive immune responses. Nat Commun 4, 2406 (2013).

38. Shokhirev, M.N. et al. A multi-scale approach reveals that NF-kappaB cRel enforces a B-cell decision to divide. Mol Syst Biol 11, 783 (2015).

39. Deenick, E.K., Gett, A.V. & Hodgkin, P.D. Stochastic model of T cell proliferation: a calculus revealing IL-2 regulation of precursor frequencies, cell cycle time, and survival. J Immunol 170, 4963–4972 (2003).

40. Hawkins, E.D. et al. Measuring lymphocyte proliferation, survival and differentiation using CFSE time-series data. Nature protocols 2, 2057–2067 (2007).

41. Hommel, M. & Hodgkin, P.D. TCR affinity promotes CD8+ T cell expansion by regulating survival. J Immunol 179, 2250–2260 (2007).

42. Bouillet, P. et al. Proapoptotic Bcl-2 relative Bim required for certain apoptotic responses, leukocyte homeostasis, and to preclude autoimmunity. Science (New York, N.Y.) 286, 1735–1738 (1999).

43. Kotschy, A. et al. The MCL1 inhibitor S63845 is tolerable and effective in diverse cancer models. Nature 538, 477–482 (2016).

44. Souers, A.J. et al. ABT-199, a potent and selective BCL-2 inhibitor, achieves antitumor activity while sparing platelets. Nat Med 19, 202–208 (2013).

45. Leverson, J.D. et al. Exploiting selective BCL-2 family inhibitors to dissect cell survival dependencies and define improved strategies for cancer therapy. Sci Transl Med 7, 279ra240 (2015).

46. Chappaz, S. et al. Homeostatic apoptosis prevents competition-induced atrophy in follicular B cells. Cell Rep 36, 109430 (2021).

47. Ke, F.F.S. et al. Embryogenesis and Adult Life in the Absence of Intrinsic Apoptosis Effectors BAX, BAK, and BOK. Cell 173, 1217–1230 e1217 (2018).

48. Lindsten, T. & Thompson, C.B. Cell death in the absence of Bax and Bak. Cell Death Differ 13, 1272–1276 (2006).

49. Wei, M.C. et al. Proapoptotic BAX and BAK: a requisite gateway to mitochondrial dysfunction and death. Science (New York, N.Y.) 292, 727–730 (2001).

50. Letai, A. et al. Distinct BH3 domains either sensitize or activate mitochondrial apoptosis, serving as prototype cancer therapeutics. Cancer Cell 2, 183–192 (2002).

51. Duke, R.C. & Cohen, J.J. IL-2 addiction: withdrawal of growth factor activates a suicide program in dependent T cells. Lymphokine Res 5, 289–299 (1986).

52. Kale, J., Osterlund, E.J. & Andrews, D.W. BCL-2 family proteins: changing partners in the dance towards death. Cell Death & Differentiation 25, 65–80 (2018).

53. Kim, H. et al. Hierarchical regulation of mitochondrion-dependent apoptosis by BCL-2 subfamilies. Nat Cell Biol 8, 1348–1358 (2006).

54. Willis, S.N. & Adams, J.M. Life in the balance: how BH3-only proteins induce apoptosis. Curr Opin Cell Biol 17, 617–625 (2005).

55. Adams, J.M. & Cory, S. The Bcl-2 protein family: arbiters of cell survival. Science (New York, N.Y.) 281, 1322–1326 (1998).

56. Korsmeyer, S.J. Bcl-2: an antidote to programmed cell death. Cancer Surv 15, 105–118 (1992).

57. Reed, J.C. Apoptosis mechanisms: implications for cancer drug discovery. Oncology (Williston Park) 18, 11–20 (2004).

58. Banjara, S., Suraweera, C.D., Hinds, M.G. & Kvansakul, M. The Bcl-2 Family: Ancient Origins, Conserved Structures, and Divergent Mechanisms. Biomolecules 10 (2020).

59. Strasser, A. & Vaux, D.L. Viewing BCL2 and cell death control from an evolutionary perspective. Cell Death & Differentiation 25, 13–20 (2018).

60. Luedtke, D.A. et al. Inhibition of Mcl-1 enhances cell death induced by the Bcl-2-selective inhibitor ABT-199 in acute myeloid leukemia cells. Signal Transduct Target Ther 2, 17012 (2017).

61. Prukova, D. et al. Cotargeting of BCL2 with Venetoclax and MCL1 with S63845 Is Synthetically Lethal In Vivo in Relapsed Mantle Cell Lymphoma. Clin Cancer Res 25, 4455–4465 (2019).

62. Horton, M.B., Hawkins, E.D., Heinzel, S. & Hodgkin, P.D. Speculations on the evolution of humoral adaptive immunity. Immunology & Cell Biology 98, 439–448 (2020).

63. Kwon, K. et al. Instructive role of the transcription factor E2A in early B lymphopoiesis and germinal center B cell development. Immunity 28, 751–762 (2008).

64. Lindsten, T. et al. The combined functions of proapoptotic Bcl-2 family members bak and bax are essential for normal development of multiple tissues. Molecular cell 6, 1389–1399 (2000).

65. Takeuchi, O. et al. Essential role of BAX,BAK in B cell homeostasis and prevention of autoimmune disease. Proc. Natl. Acad. Sci. U. S. A. 102, 11272–11277 (2005).

66. Storn, R. & Price, K. Differential evolution - A simple and efficient heuristic for global optimization over continuous spaces. J Global Optim 11, 341–359 (1997).

67. Parks, D.R., Roederer, M. & Moore, W.A. A new “Logicle” display method avoids deceptive effects of logarithmic scaling for low signals and compensated data. Cytometry A 69, 541–551 (2006).

68. Castillo-Hair, S.M. et al. FlowCal: A User-Friendly, Open Source Software Tool for Automatically Converting Flow Cytometry Data from Arbitrary to Calibrated Units. ACS Synth Biol 5, 774–780 (2016).

