## Supplementary Materials for "A quantitative molecular model of the lymphocyte death timer"

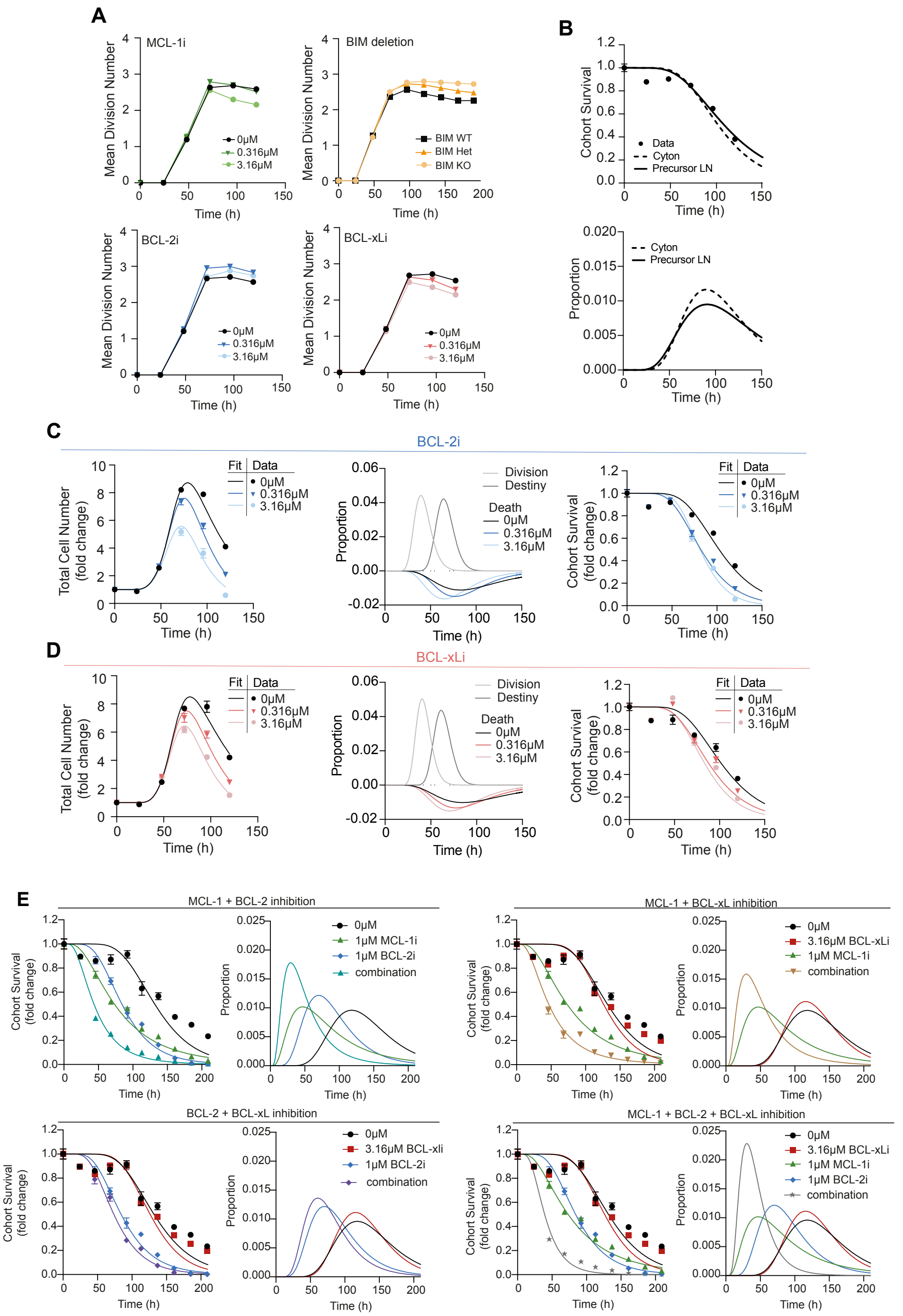

**Supplementary Figure 1. Manipulation of BCL-2 family protein availability and effects on CpG responses.**

**(a)** Mean division number analysis of CTV labelled B cells stimulated with CpG and treated with BCL-2i, MCL-1i and BCL-xLi 24h after activation or following BIM deletion. **(b)** Cohort survival curve and death time distribution comparison using Cyton modelling or direct lognormal fitting of measured data. **(c-e)** Cohort survival data with fitted lognormal survival curves and death time distributions of CpG stimulated B cells in response to single or combination BH3 mimetic treatment as indicated. Data represented as mean ± SEM.

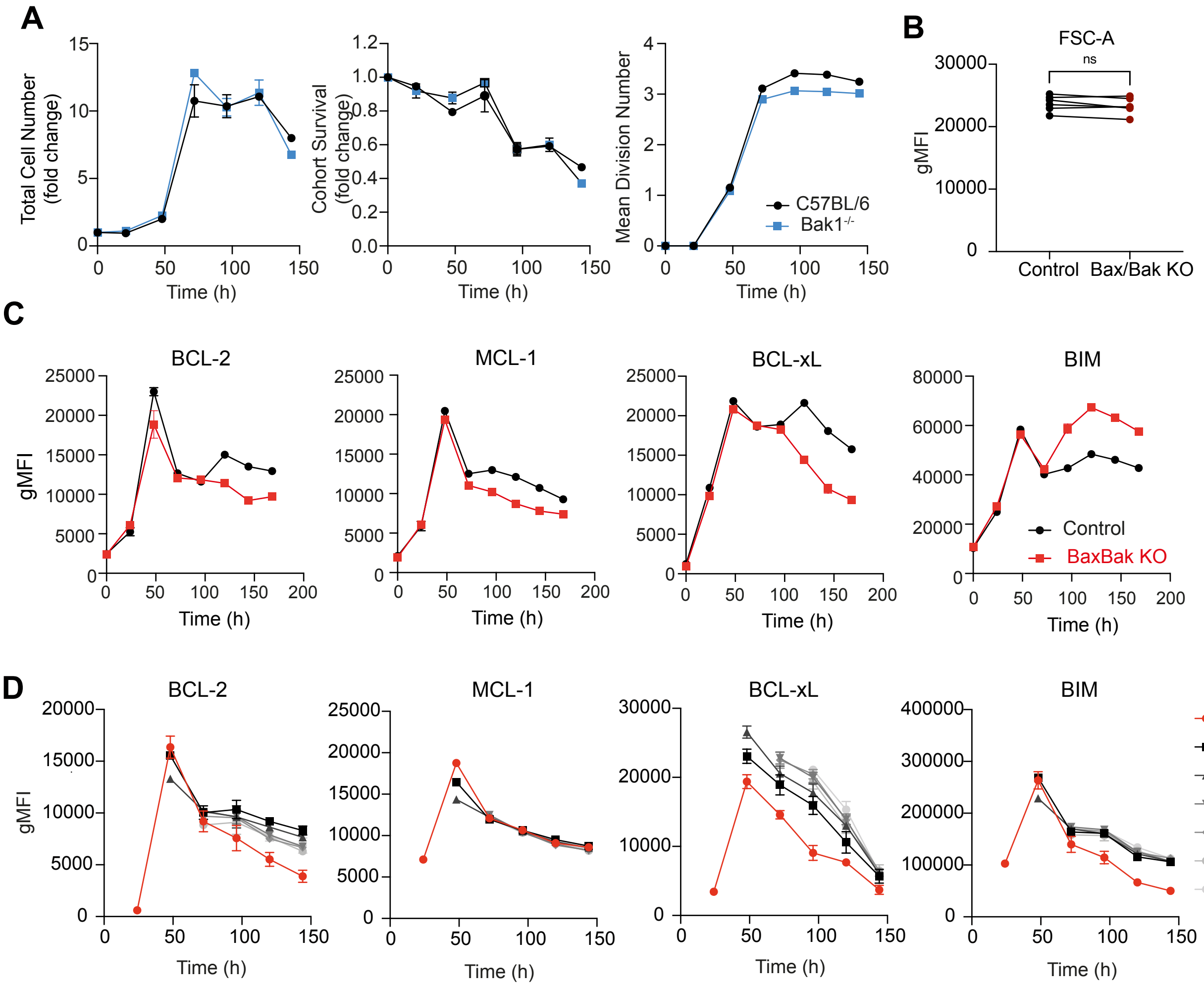


**Supplementary Figure 2. Response of BAK-deficient B cells to CpG stimulation and the impact of cell division on BCL-2 family expression in Bax/Bak KO B cells.**

**(a)** A comparison of the response of B cells from *Bak1*$^{-/-}$ and wildtype C57BL/6 mice (total cell number, cohort survival and mean division number over time) to CpG stimulation. Data represented as mean ± SEM. **(b)** Cell size as indicated by forward scatter area (FSC-A) of naïve B cells isolated from either control or BaxBak KO mice. A paired t-test was performed on paired control and BaxBak KO data from six independent experiments (NS). Purified, CTV labelled Bax/Bak KO B cells were stimulated with CpG and expression of BCL-2, MCL-1, BCL-xL and BIM was quantified **(c)** on the population level and **(d)** per division over time. Data represented as mean ± SEM.

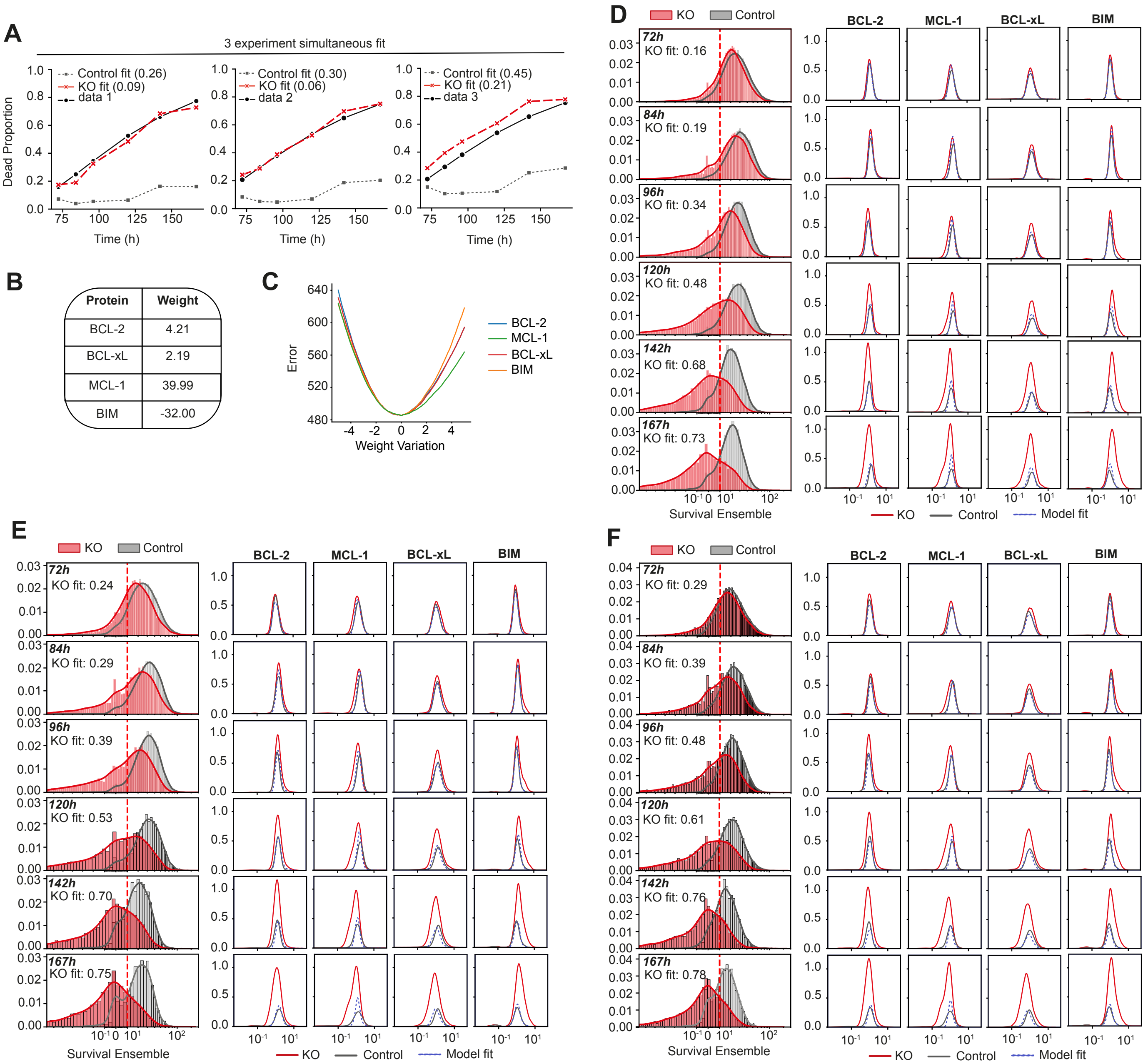

**Supplementary Figure 3. Simultaneous Ensemble-Threshold model (linear sum) analysis of multiple data sets.** Simultaneous ET model fitting was performed for the three data sets. **(a)** Death curve predictions, **(b)** the fitted protein weights and **(c)** the sensitivity of fitting to changes in protein weights are plotted. ET model survival ensembles and protein expression predictions for **(d)** Data 1, **(e)** Data 2 and **(f)** Data 3 were calculated.

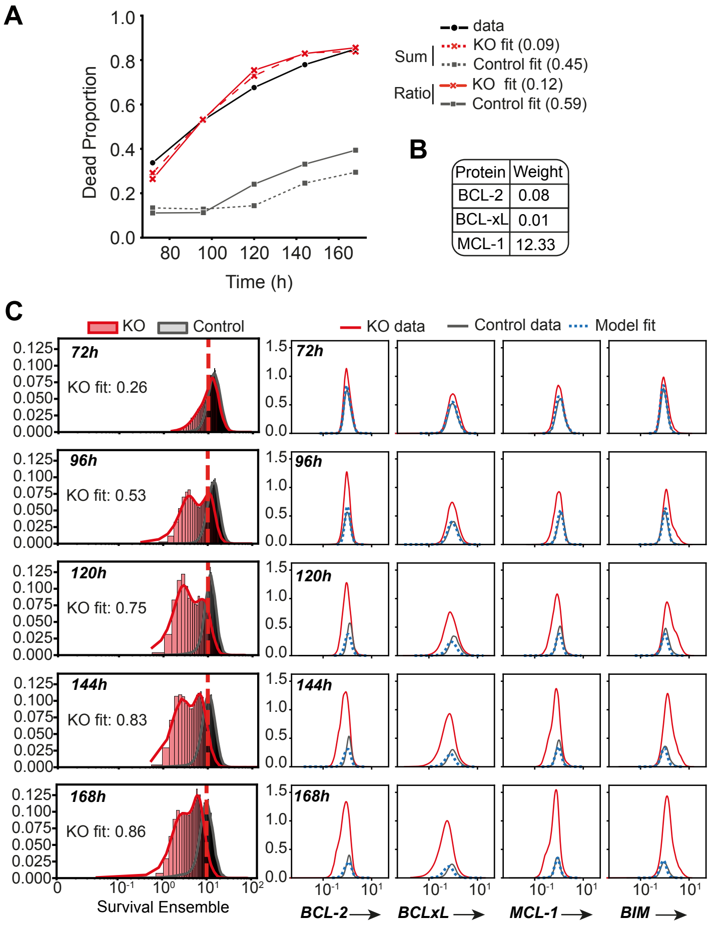


**Supplementary Figure 4. Ensemble-Threshold model (Ratio) analysis.**

Ratio ET model fitting was performed for the B cell survival and BCL-2 family expression data. **(a)** Death curve predictions, including comparison of death curve predictions using ET model (linear sum), as in Figure 3, **(b)** the fitted protein weights of the ratio model and **(c)** ET model survival ensembles and protein expression predictions were calculated.

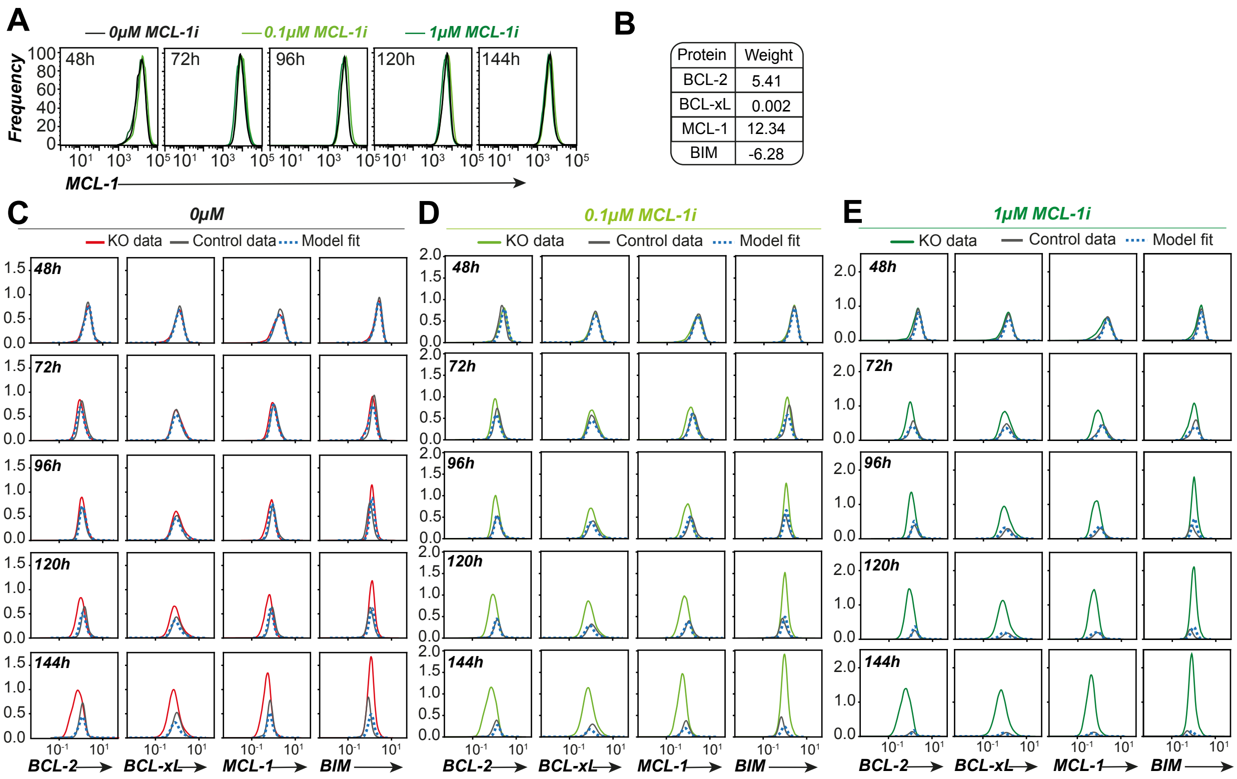


**Supplementary Figure 5. The effect of MCL-1i treatment on protein expression, survival, and ET model parameters.** Purified, CTV labelled B cells were stimulated with CpG and treated 24h with MCL-1i as indicated. (**a**)The effect of MCL-1i treatment on MCL-1 protein expression was assessed in Bax/Bak KO cells**.** Data in A are representative plots from triplicate samples. Undrugged experimental data was fitted using ET death model (Figure 4c), to calculate (**b**) protein weights and **(c)** protein histogram predictions were determined. Protein histogram predictions following ET model fitting and incorporation of IFs for **(d)** 0.1uM MCL1i or **(e)** 1uM MCL1i data.

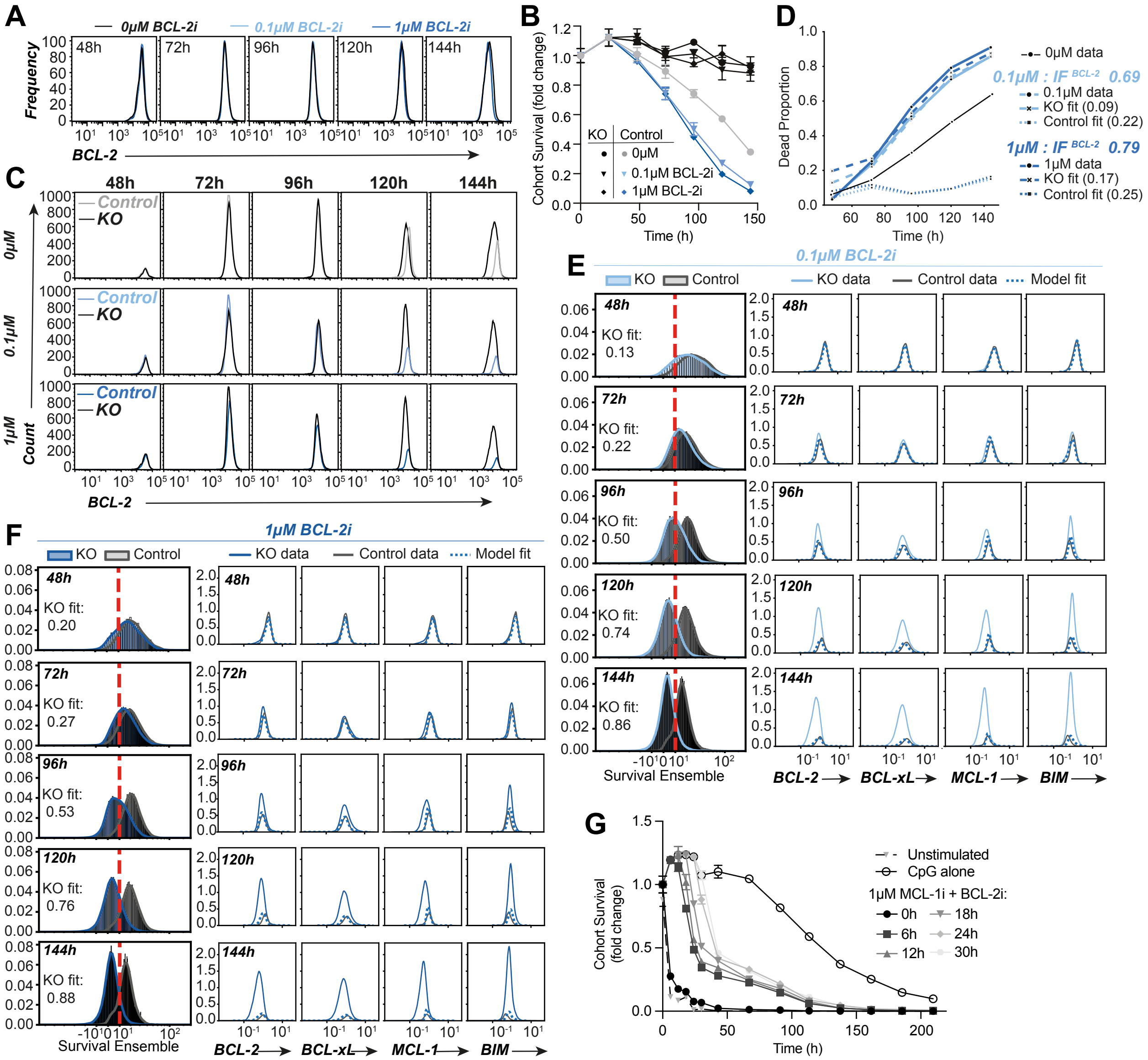

**Supplementary Figure 6. The effect of BCL-2i treatment on protein expression, survival, and ET model parameters.**

Purified, CTV labelled B cells were stimulated with CpG and treated at 24h with BCL-2i as indicated. (**a**)The effect of BCL-2i treatment on BCL-2 protein expression was assessed in Bax/Bak KO cells**.** Data in A are representative plots from triplicate samples. (**b**) Cohort Survival and **(c)** BCL-2 expression were determined over time. **(d)** ET model death curve fits and inhibition factors following BCL-2i treatment and ET model protein expression predictions for **(e)** 0.1µM BCL2i or **(f)** 1µM BCL2i. **(g)** Effect on cell survival where 1µM MCL-1i and BCL-2i combination treatment was added at various times post CpG activation. Data are shown as mean ± SEM.

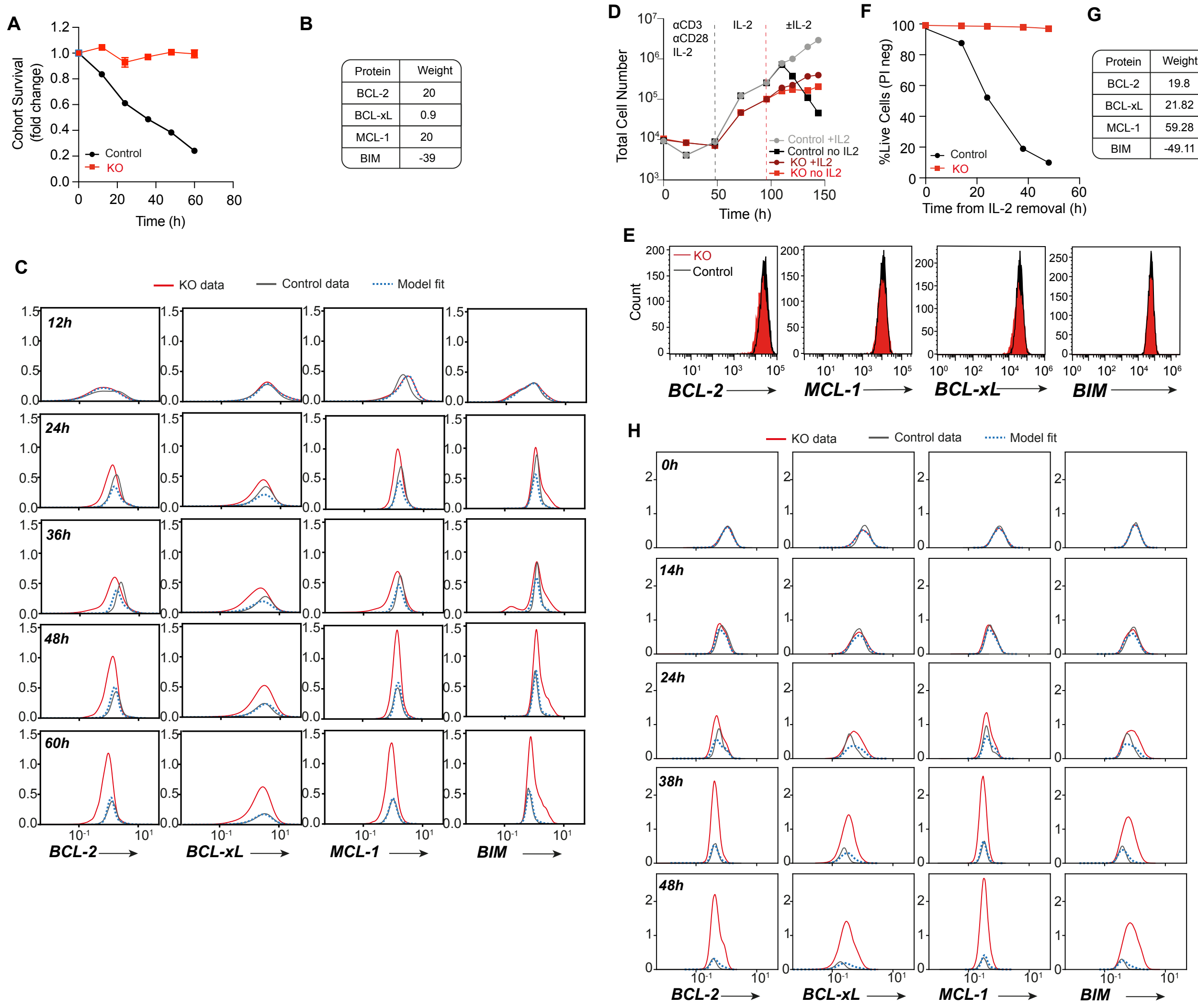


**Supplementary Figure 7. ET cell death model analysis of unstimulated B cells and $CD4^+$ T cells following IL-2 withdrawal**

**(a)** Cohort survival of unstimulated control or Bax/Bak KO B cells were analyzed over time. Data represented as mean ± SEM. ET model analysis of unstimulated control and Bax/Bak KO B cells was performed and (**b**) fitted protein weights and (**h**) ET model fits of BCL-2 family protein expression overlaid with observed data. **(c)** Total control or Bax/Bak KO $CD4^+$ T cell numbers were analyzed after activation with αCD3/αCD28 and IL-2. After subsequent culture in saturating levels of IL-2, cells were split and cultured in the presence or absence of IL-2 as indicated from 96h. Data represented as mean ± SEM. (**e**) Expression of BCL-2 family proteins in control and Bax/Bak KO $CD4^+$ T cells at 96h, the point of IL-2 withdrawal. (**f**) $CD4^+$ T cell survival, as indicated by propidium iodide staining, was determined after IL-2 withdrawal. ET model analysis of activated control and Bax/Bak KO $CD4^+$T cells following IL-2 withdrawal was performed and (**g**) fitted protein weights and (**h**) ET model fits of BCL-2 family protein expression overlaid with observed control and Bax/Bak KO data. Data are representative of two independent experiments.
